# Model-based projections of Zika virus infections in childbearing women in the Americas

**DOI:** 10.1101/039610

**Authors:** T. Alex Perkins, Amir S. Siraj, Corrine Warren Ruktanonchai, Moritz U.G. Kraemer, Andrew J. Tatem

## Abstract

Zika virus is a mosquito-borne pathogen that is rapidly spreading across the Americas. Due to associations between Zika virus infection and a range of fetal maladies^1,2^, the epidemic trajectory of this viral infection poses a significant concern for the nearly 15 million children born in the Americas each year. Ascertaining the portion of this population that is truly at risk is an important priority. One recent estimate^3^ suggested that 5.42 million childbearing women live in areas of the Americas that are suitable for Zika occurrence. To improve on that estimate, which did not take into account the protective effects of herd immunity, we developed a new approach that combines classic results from epidemiological theory with seroprevalence data and highly spatially resolved data about drivers of transmission to make location-specific projections of epidemic attack rates. Our results suggest that 1.65 (1.45–2.06) million childbearing women and 93.4 (81.6–117.1) million people in total could become infected before the first wave of the epidemic concludes. Based on current estimates of rates of adverse fetal outcomes among infected women^2,4,5^, these results suggest that tens of thousands of pregnancies could be negatively impacted by the first wave of the epidemic. These projections constitute a revised upper limit of populations at risk in the current Zika epidemic, and our approach offers a new way to make rapid assessments of the threat posed by emerging infectious diseases more generally.

---

On February 1, 2016, the World Health Organization (WHO) designated the ongoing Zika virus epidemic in the Americas as a Public Health Emergency of International Concern (PHEIC), defined as an “extraordinary event” that “potentially require[s] a coordinated international response”^6^. This declaration acknowledges the high potential for Zika to establish across the Americas given that its dominant vector, the *Aedes aegypti* mosquito, is endophilic and occupies an exceptionally broad range^7^. Concern underlying this rare WHO declaration also stems from an association between Zika virus infection in pregnant women and a range of adverse fetal outcomes^2^, most notably congenital microcephaly^1^. As of April 28, 2016, there were 1,286 confirmed cases of microcephaly associated with Zika virus infection in five countries^8^, and there is widespread concern that these numbers could increase further as the virus continues to spread across the Americas^9^.

A number of uncertainties surround the future of the Zika epidemic in the Americas, particularly questions about how many women may be at risk of having children with congenital microcephaly and other adverse outcomes associated with Zika virus infection^10^. Of women who become infected with Zika virus during a vulnerable stage of their pregnancy, evidence is emerging that 1-13% may go on to develop congenital microcephaly^2,4,5^. However, the number of women who become infected with Zika virus during that timeframe is difficult to ascertain. One recent study^3^ estimated that 5.42 million births occurred in 2015 in regions of the Americas with “suitability” for Zika “occurrence.” Such estimates come with many caveats though, as they rely on a relatively limited number of reported cases and apply a method based on equilibrium assumptions to a situation involving active range expansion^11^. Most importantly, the estimate of 5.42 million births^3^ reflects the total population within a demarcated area and does not take into account that large fractions of the populations in those areas may remain uninfected due to herd immunity and other factors_12,13_.

To quantify the potential magnitude of the ongoing Zika epidemic in terms of people who realistically might become infected, we formulated and applied a method for projecting location-specific epidemic attack rates on highly spatially resolved human demographic projections^14^. The central concept behind our approach is that of the “first-wave” epidemic. Zika and other mosquito-borne viruses have been known to exhibit explosive outbreaks, infecting as much as 75% of a population in a single year^15^. Classical epidemiological theory predicts that some proportion of a population will remain uninfected during an epidemic, because herd immunity eventually causes the epidemic to burn out^12^. A related prediction of this theory is that the proportion infected prior to epidemic burnout (i.e., the epidemic attack rate) has a one-to-one relationship with the basic reproduction number, *R*_0_^13^. The latter quantity has a well-known mechanistic formulation for mosquito-borne pathogens^16^ that accommodates the effects of environmental drivers on transmission_17,18_. For example, the incubation periods of dengue viruses in the *Ae. aegypti* mosquitoes that transmit Zika virus have an empirically derived relationship with temperature^18^, which can in turn be used to inform calculations of *R*_0_. Together with similar relationships for other transmission parameters, it is possible to characterize *R*_0_, a fundamental measure of transmission potential, as a function of local environmental conditions.

We leveraged these classic results from epidemiological theory to first perform highly spatially resolved calculations of *R*_0_ and then to translate those calculations into location-specific projections of first-wave epidemic attack rates (Fig. 1a). Because Zika-specific values of transmission parameters are largely unknown at present but may be well approximated by dengue-specific values^19^, we used some parameter values for dengue virus in our *R*_0_ calculations. We furthermore calibrated our attack rate projections to match empirically estimated attack rates from 12 chikungunya epidemics and one Zika epidemic in naïve populations (Extended Data Table 1). This step afforded us the flexibility to enhance the realism of the model with respect to firmly established but poorly quantified associations between human-mosquito contact and economic prosperity^20^. In doing so, one departure from the classic relationship between *R*_0_ and attack rate that we made was to rescale *R*_0_ by an exponent *α* ∈ (0,1] to allow for better correspondence with observed attack rates. Although there is no theoretical justification for this or any other particular scaling relationship, it is consistent with theoretical expectations^21^ that attack rates should be lower in populations with equal *R*_0_ values but more heterogeneous contact patterns, which are typical for transmission by *Ae. aegypti*^*22*^. To provide a point of reference for our model-based approach, we also fitted a statistical description of the 13 seroprevalence estimates as a function of the environmental drivers that we considered. For both approaches, we applied their respective location-specific attack rate projections to demographic projections on a 5x5 km grid across Latin America and the Caribbean to obtain expected numbers of infections in the overall population and among childbearing women in particular (Fig. 1c). All such calculations were performed for 1,000 Monte Carlo samples of model parameters.

**Figure 1.**
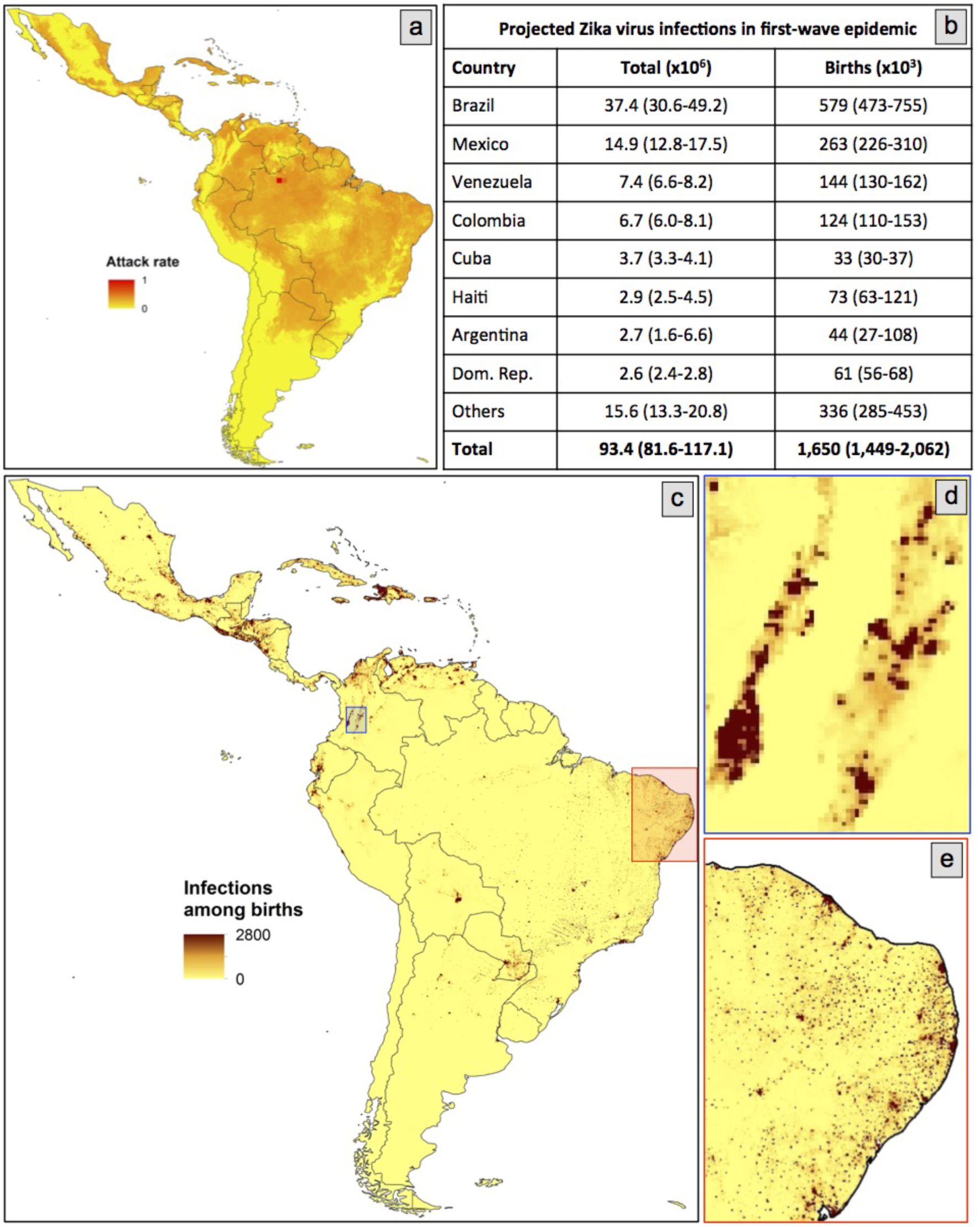
Spatial projections of location-specific epidemic attack rates (a) that combine with demographic projections^14^ to yield projections of total numbers of Zika infections in all people and in childbearing women, which can either be summarized by country (b) or on a map (c). More detailed spatial projections of infections among childbearing women are shown for two areas: Cali, Colombia (d) and Recife, Brazil (e).

In total, our median projection suggests that as many as 93.4 (range: 81.6-117.1) million people in Latin America and the Caribbean could become infected during the first wave of the epidemic (Fig. 1b). To place this number into context, we refer to an estimate^23^ that 53.8 (40.0–71.8) million dengue infections occurred in this region in 2010 alone. Our projections of nearly double this number for Zika are not surprising, given that there is extensive immunity to dengue but not Zika in this region and given that it would likely take longer than a year for the first wave of the epidemic to conclude in all locations within this region. At the country level, we project that Brazil will have the largest total number of infections by more than double that of any other country, due to a combination of its size and suitability for transmission. Island countries in the Caribbean are projected to experience the highest nationally averaged attack rates, with 7 of the highest 10 values projected for countries including Aruba, Haiti, and Cuba. This projection is consistent with a frequent history of arbovirus outbreaks on islands^24^ and may owe to the uniformity of environmental conditions on the portions of islands where people tend to live. In more heterogeneous regions, the 5x5 km spatial resolution of our maps allows for nuanced projections for areas of interest to local stakeholders (Fig. 1d,e).

Among childbearing women, our median projection suggests that there could be as many as 1.65 (range: 1.45–2.06) million infections in Latin America and the Caribbean before the first wave of the epidemic concludes (Fig. 1b). Assuming that birth rates are temporally constant, our projections are robust to uncertainty about the timing of local epidemics and the timeframe of the first wave of the epidemic, because they are based on cumulative proportions infected. These projections can also be used to postulate numbers at risk of microcephaly by multiplying them by the fraction of a year in which a pregnant woman is susceptible to developing microcephaly (e.g., multiply by 1/4 in the case of first-trimester susceptibility). We also note that there were some discrepancies in our projections in terms of the rank order of countries experiencing the most infections among childbearing women versus the population as a whole. In particular, Cuba was 5^th^ in terms of projected infections in the overall population but 12^th^ in terms of infections among childbearing women due to its low birth rate compared to other countries in the Americas^25^. Such discrepancies are also likely to exist subnationally^26^, and their elucidation should be a priority for future work.

By accounting for uncertainty distributions for each of the key drivers of our model (Fig. 2a–e), we found that uncertainty distributions for infections across the region as a whole and by country were often multimodal (Fig. 2f–o) due to uncertainty in the shape of the relationship between mosquito-human contact and the local economic index that we considered (Fig. 2d). Summing our projections across Latin America and the Caribbean revealed variation that was modest, in the sense that none of our 1,000 Monte Carlo samples resulted in fewer than 81 million infections overall and 1.4 million among childbearing women (Fig. 2f,k). There are many reasons that even these numbers could be overestimates though. Our projections are conditioned on a local epidemic taking place in each 5x5 km grid cell in the region, which is unlikely to happen given dispersal limitation, stochastic fadeout, geographic mismatches in seasonality, and other factors. Therefore, it is most appropriate to interpret our projections as either a plausible worst-case scenario or an expectation of local epidemic size conditional on there being a local epidemic in the first place.

**Figure 2.**
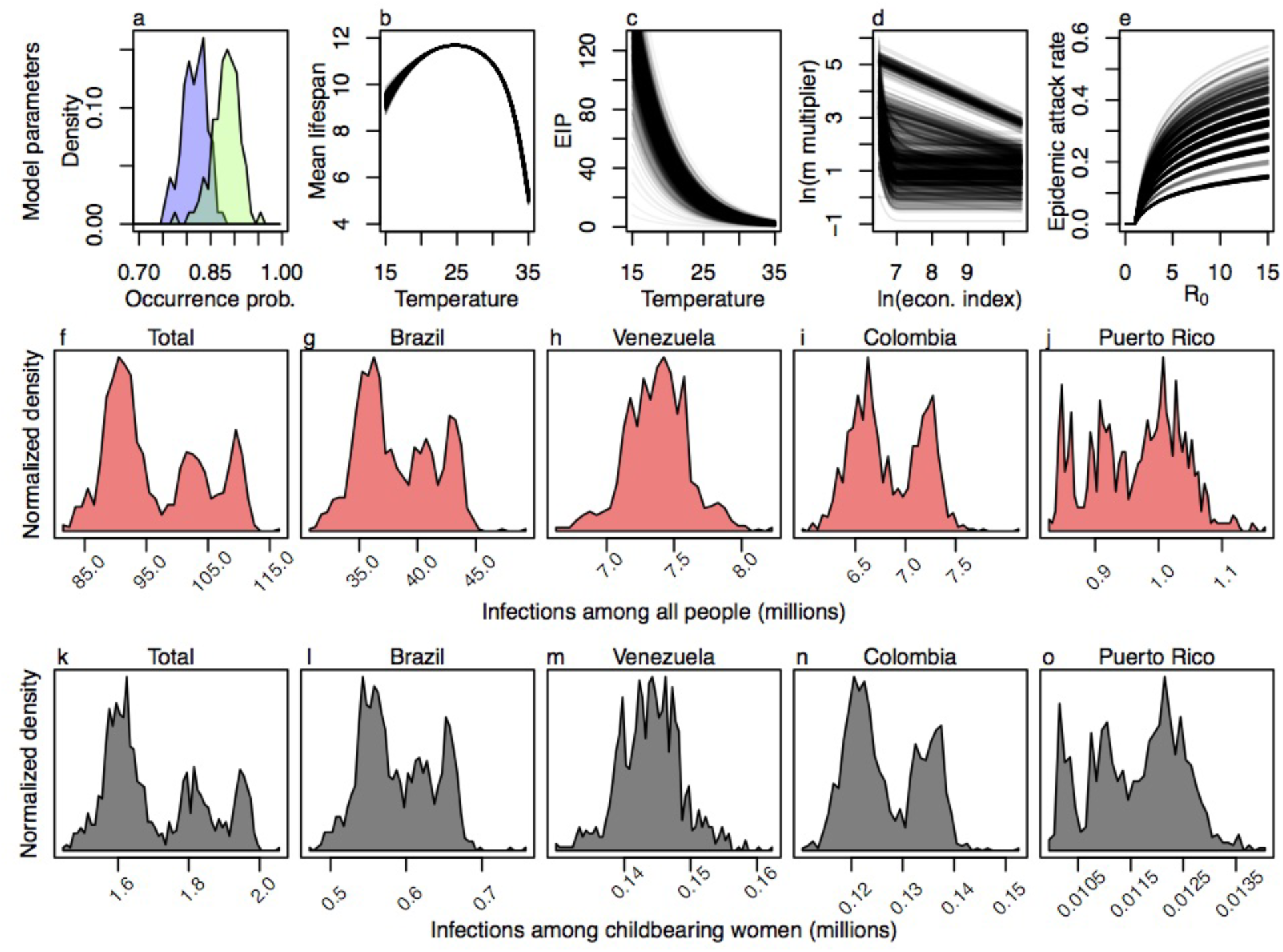
Monte Carlo samples from the uncertainty distributions surrounding each of the key drivers in the model (a-e) and uncertainty distributions for projected numbers of infections among all individuals (f-j) and among childbearing women (k-o) in different areas. All panels reflect the full range of uncertainty considered in 1,000 Monte Carlo samples. Panel a shows posterior distributions of mosquito occurrence probabilities for two example 5x5 km grid cells.

Although our approach was very much rooted in mechanistic models from epidemiological theory, two critical steps in our method involved fitting curves to describe theoretically motivated but heretofore unknown relationships: an association between mosquito-human contact and economic prosperity (Fig. 2d), and a scaling relationship between *R*_0_ and attack rates (Fig. 2e). Allowing these relationships to be informed by local seroprevalence estimates (Extended Data Table 1) left open the question of the extent to which our projections were informed by the mechanistic assumptions of the model versus statistical fits to the seroprevalence estimates that we used. On the one hand, an alternative statistical approach accounted for much more variation in seroprevalence estimates (*R*^2^=0.89) than did the model-based approach (*R*^2^=0.32). On the other hand, the statistical approach offered a dichotomous set of projections about numbers of infections outside the context of the data to which it was fitted: either everyone will become infected or very few people will (Fig. 3). Relationships between attack rates and predictor variables inferred by the statistical approach (Fig. 4d–i) were also implausible: a narrow temperature range in which attack rates increase sharply towards 100% (Fig. 4d–h), and a reversal of economic effects whereby wealthy populations experience higher attack rates than poor populations when mosquito occurrence probabilities are high (Fig. 4f,i).By contrast, the model-based approach yielded more moderate attack rate projections overall (Fig. 2f vs 3a) in which temperature, economic prosperity, and mosquito occurrence probability all had plausible relationships with attack rates (Fig. 4a–c).

**Figure 3.**
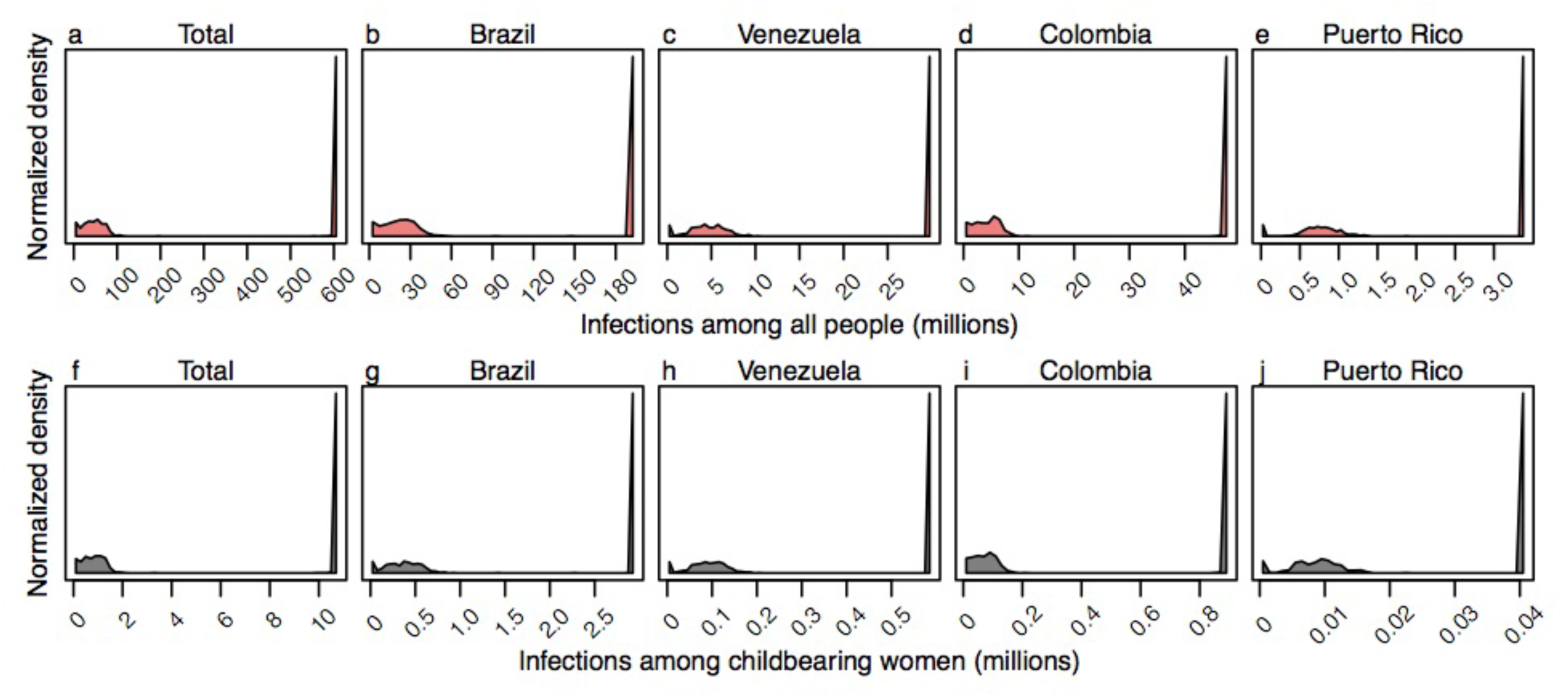
Projected numbers of infections among all individuals (a-e) and among childbearing women (f-j) for 1,000 Monte Carlo samples from the uncertainty distribution around parameters of the statistical model that was fitted to seroprevalence estimates from Extended Data Table 1.

**Figure 4.**
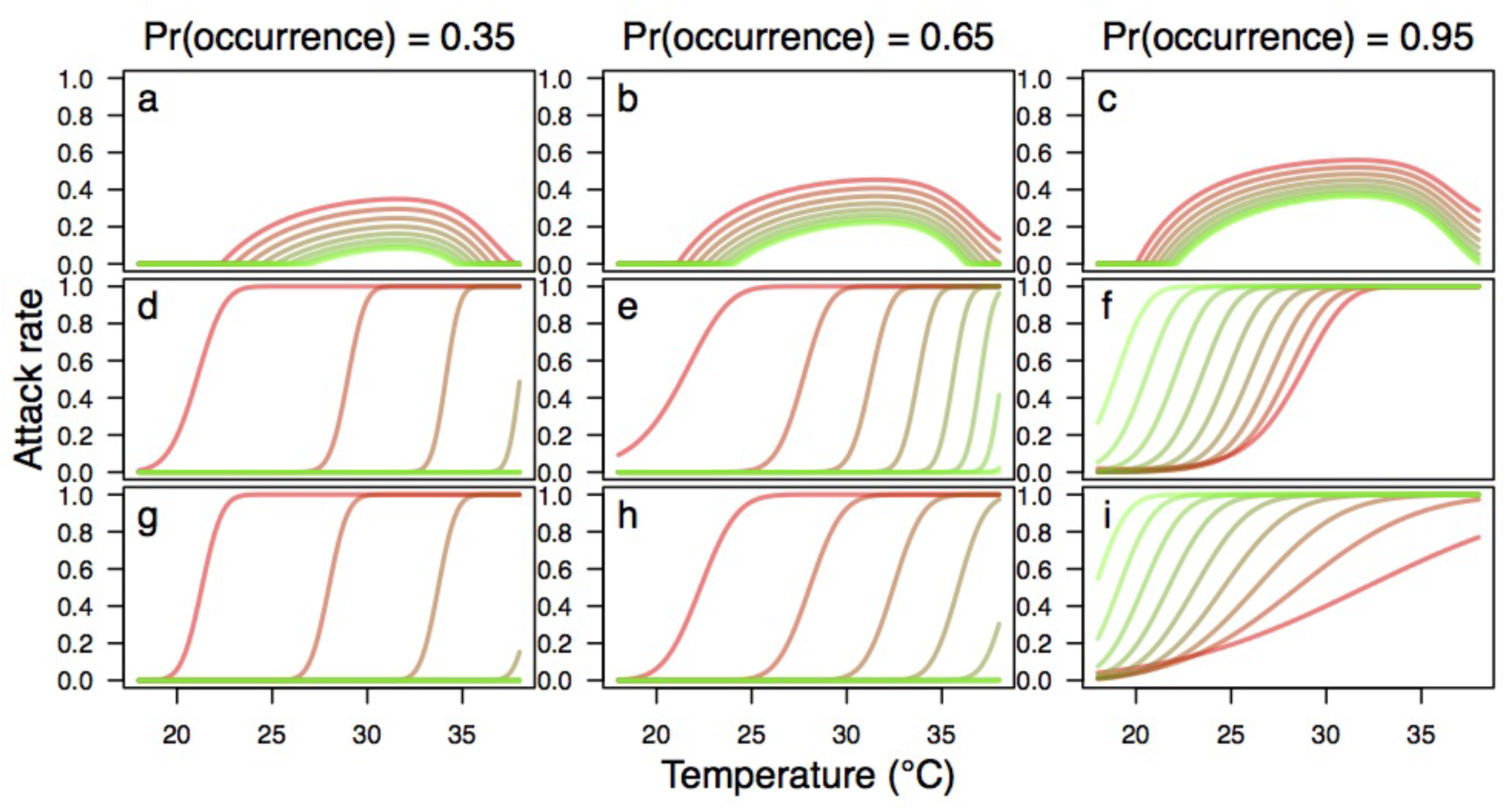
Relationships between temperature (x-axis), economic index (red = low, green = high), mosquito occurrence probabilities (columns), and projected epidemic attack rates (y-axis). These relationships are shown for the model-based approach (a-c), the full statistical approach (d-f), and the version of the statistical approach chosen by stepwise model selection (g-i).

In conclusion, our model-based approach offers a unique way to leverage a variety of spatially detailed data products ^7,14,27,28^ to make a priori projections of attack rates and infections that could be experienced in the first wave of the ongoing Zika epidemic. Projections such as these have an important role to play in the early stages of an epidemic, when planning for surveillance and outbreak response is actively underway both internationally and locally^9^. At the same time, it is important for consumers of this information to be aware of uncertainties in these and other projections, which often exceed the amount of uncertainty that can be identified a priori^29^. Likewise, following up on these projections in the aftermath of the epidemic—by comparing against projections made with alternative models and additional serological surveys^30^—will provide an exceptional opportunity to enhance capabilities to anticipate the severity of future epidemic threats.

## METHODS

### Data sources and processing

#### Human demography

To estimate the annual numbers of pregnancies per 1x1 km grid cell in 2015, methods developed by the WorldPop project (www.worldpop.org^25,31^) were adapted for the Americas region. Highresolution estimates of population counts per 100x100 m grid cell for 2015 were recently constructed for Latin American, Asian, and African countries^14,32^. With consistent subnational data on sex and age structures, as well as subnational age-specific fertility rate data across the Americas currently unavailable for fully replicating the approaches of Tatem et al.^31^, national level adjustments were made to construct pregnancy and birth counts. Data on estimated total numbers of births^33^ and pregnancies^31^ occurring annually in 2012 were assembled for all Latin American study countries, as well as births in 2015^33^. As no 2015 pregnancy estimates existed at the time of writing, the ratios of births to pregnancies for each country in the Americas were calculated using 2011 and 2012 estimates, and these were then applied to the 2015 births numbers to obtain 2015 estimates of annual pregnancy numbers per-country. This made the assumption that per-country births-to-pregnancies ratios remained the same in 2015 as they were in 2011 and 2012. The 100x100 m gridded population totals were aggregated to 1x1 km spatial resolution, and the per-country totals were linearly adjusted to match the 2015 pregnancy estimates.

#### Temperature

We used interpolated meteorological station temperature data from the 1950-2000 period at 5x5 km spatial resolution, processed to create climatological monthly averages that represent “typical” conditions (www.worldclim.org^27^).

#### *Aedes aegypti* occurrence probability

To predict the likely distribution of *Aedes aegypti* mosquitoes, Kraemer et al.^7^ generated highresolution occurrence probability surfaces based on a species distribution modeling approach^11^. More specifically, a boosted regression tree model was applied using a comprehensive set of known occurrences (*n* = 19,930) of *Ae. aegypti* and a set of environmental predictors known to influence the distribution of the species^7^. Covariates included a temperature suitability index^17^, contemporary mean and range maps of the Enhanced Vegetation Index and precipitation^34^, and an urbanization index from the Global Rural Urban Mapping Project. We used a set of 100 spatial layers sampled from the posterior distribution estimated by Kraemer et al.^7^.

#### Economic index

To account for socio-economic differences among populations residing in different regions, we used one-degree resolution gridded estimates of purchasing power parity (PPP) in U.S. Dollars from 2005 adjusted for inflation (G-Econ)^28^. When we encountered missing values, we imputed values in one of two ways. Grid cells in small island countries with data missing for the entire country were uniformly filled with population-adjusted PPP figures obtained from the U.S. CIA World Factbook^35^. Missing values in continental grid cells were imputed with the mean of the surrounding eight grid cell values. Once we obtained a complete PPP grid layer at one-degree resolution, we resampled the layer to a resolution of 5x5 km to match the resolution of gridded layers for human demography, temperature, and *Ae. aegypti* occurrence probability.

#### Seroprevalence estimates

To calibrate our model, we identified published estimates of seroprevlance that were relevant to the context of our study (Extended Data Table 1). Specifically, we sought estimates of seroprevalence to either Zika or chikungunya viruses in populations that were presumably naïve prior to an outbreak. Thus, we excluded some seroprevalence estimates that were obtained from endemic populations. We also excluded estimates from small islands—namely, Reunion and Grande Comore—for which it was clear that gridded temperature data were unrealistically low due to steep elevational gradients and other features of island geography. Although the focus of our analysis was on Latin America and the Caribbean, we were not able to exclude locations on the basis of location given that only 2/13 came from the focal region. Appropriately however, a number of the seroprevalence estimates we obtained pertained specifically to pregnant women, although there did not appear to be differences in the seroprevalence of pregnant women and the population at large, at least in the context of a naïve population following an outbreak^36^.

### Calculation of derived quantities

#### Mosquito abundance

Occurrence probabilities can be translated into proxies for abundance provided that an assumption is made about how abundance is distributed as a random variable^37^. Assuming that mosquito abundance is distributed as a Poisson random variable, the probability that there is at least one mosquito present in a given location is 1 — exp(—*λ*), where *λ* is the expected abundance of mosquitoes. Inverting this relationship, we obtained an estimate *λ* = — ln(1 — occurrence probability) of expected mosquito abundance under the Poisson model and used this as a proxy for mosquito abundance in our calculations.

#### Mosquito-human ratios

The estimates of mosquito occurrence probability that we used incorporated a number of environmental variables^7^. They did not account for factors that modulate contact between mosquitoes and humans, however. Due in part to economic differences, factors such as air conditioning and piped water can drastically limit mosquito-human contact and virus transmission, even when mosquitoes are abundant^20^. We accounted for the effect of economic differences between locations by multiplying our proxy for mosquito abundance *λ* by a multiplication factor modeled with a function of the gross cell product economic index that we fitted to match our attack rate projections with published seroprevalence estimates. We fitted this function describing the multiplication factor using a shape constrained additive model (SCAM^38^), because relationships between mosquito-human ratios and economic indices should be monotonically decreasing, nonlinear, and without a predictable functional form. The response variable to which we fitted this relationship was a set of scalar multiples of expected mosquito abundances that would result in a perfect correspondence between attack rates derived from our model and published seroprevalence estimates.

#### Basic reproduction number *R*_0_

We calculated the basic reproduction number *R*_0_ according to its classic Ross-Macdonald formulation and as a function of temperature *T*,

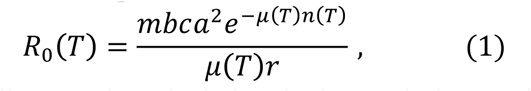

with adult mosquito mortality *μ* and extrinsic incubation period *n* specified as functions of temperature. Because temperature values were available for each location on a monthly basis, we computed monthly values of *R*_0_ for each location and then used the mean of the highest six monthly values of *R*_0_ as a singular estimate of *R*_0_ for each location. This approach was broadly consistent with the way in which a temperature suitability index was used to inform mosquito occurrence probabilities by Kraemer et al.^7^.

For mosquito mortality, we used the temperature‐ and age-dependent model of Brady et al.^39^, to which we added an additional force of extrinsic mortality (0.025 d^−1^) to match an overall daily mortality value of 0.115 estimated in a mark-release-recapture experiment carried out under temperatures ranging 20-34 °C ^40^. We then computed the mean of the age‐ and temperature-dependent lifespan distribution as a function of temperature to inform *μ*(*T*). For the relationship between temperature and mean duration of the extrinsic incubation period, we used the temperature-dependent exponential rate estimated by Chan and Johansson^18^. The ratio of mosquitoes to humans, *m*, was quantified using a combination of occurrence probabilities and the gross cell product economic index, as described in the previous two sections. Parameters that did not depend on temperature were set at the following values according to published estimates for *Ae. aegypti* and dengue virus: mosquito-to-human transmission probability, *b* = 0.4 ^41^; human-to-mosquito transmission probability times number of days of human infectiousness, *c* / *r* = 3.5 ^42^; and mosquito biting rate, *a* = 0.67 ^43^. Although there is uncertainty around these parameter values, any such uncertainty was effectively subsumed by fitting *m* to seroprevalence data given that *bca*^2^/*r* entered *R*_*0*_ as a constant.

#### Attack rates under a model-based formulation

Under a susceptible-infected-recovered (SIR) transmission model, there is a one-to-one relationship between *R*_0_ and final epidemic size, which is equivalent to the attack rate over the course of an epidemic^13^. Intuitively, the final epidemic size is reached once herd immunity is sufficient to limit contacts between infectious and susceptible individuals to the extent necessary to reduce the pathogen’s force of infection to zero. There is no explicit solution for final epidemic size as a function of model parameters, but it can be calculated numerically by obtaining an implicit solution of *S*_∞_ = *e*^−*R*_0_(1–*S*_∞_)^ for *S*_∞_), which is the proportion remaining susceptible after the epidemic has burned out^13^. Under the assumptions of the SIR model, the attack rate over the course of an epidemic is *AR* = 1 – *S*_∞_.

To apply this theoretical insight to Zika or other mosquito-borne pathogens, several limiting assumptions of the SIR model must first be reconciled. One such assumption is that individuals become infectious immediately upon becoming infected and remain infectious for an exponentially distributed period of time^44^; mosquito-borne pathogens such as Zika virus are instead characterized by a distinct lag between human and mosquito infection^45^. Despite this discrepancy between assumptions of the SIR model and the reality of many pathogen systems, mathematical analyses^46^ have shown that final epidemic size is insensitive to details about the shape of the distribution that characterizes the time period between successive cases (i.e., the generation interval).

Another limiting assumption of the SIR model is that of homogeneous encounters between people and mosquitoes^44^, which are understood to be extensive for mosquito-borne diseases^22^. Mathematical analyses^21^ in this case show that a seemingly infinite complexity of relationships between *R*_0_ and final epidemic size are possible in a heterogeneous system. As a general rule, however, final epidemic size in a system with contact heterogeneity and proportional mixing is expected to be strictly less than the final epidemic size in an otherwise equivalent system with homogeneous contacts^21^. How the ratio of these final epidemic sizes scales as a function of *R*_0_ depends entirely on the details of a given system and has so far not been generalized mathematically.

To capture the potentially very strong effects of heterogeneity in reducing final epidemic size in populations subject to Zika epidemics, we scaled final epidemic size by substituting 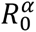 for *R*_0_ in the SIR-based final epidemic size formula given some constant *α* ∈ (0,1]. Although there is no theoretical justification for this or any other choice of how to scale *R*_0_ and *AR* in the presence of contact heterogeneity, the choice we made has the following desirable properties: (1) it implies that *AR* → 1 as *R*_0_ → ∞; (2) it leads to the function *AR*(*R*_0_) having a more gradual slope and thereby allowing for intermediate attack rates to be more common than they would be otherwise; and (3) it preserves the property that *AR* = 0 for *R*_0_ < 1. At the same time, this and possible alternative formulations are limited by a general lack of understanding about the relationship between *R*_0_ and *AR* in heterogeneous systems, relationships that may furthermore be heterogeneous themselves across different areas^47^.

To estimate *α*, we performed the following procedure for candidate values of *α* between 0.01 and 1 in increments of 0.01: (1) calculate *R*_0_ according to eqn. 1 and assuming *m* = 1 for each of the 13 sites from which seroprevalence estimates were derived; (2) use those *R*_0_ values to calculate *AR* values for each of those sites based on the classic SIR formulation; (3) calculate what multiplication factor of *R*_0_ would be necessary for *AR* to match the empirical seroprevalence estimate; (4) fit a SCAM model of the economic index to the multiplication factors; and (5) use the fitted SCAM values to recalculate *R*_0_ and then *AR* for each site. Next, we calculated the sum of squares between the final predicted *AR* values associated with each *α* and the empirical seroprevalence estimates, and we then selected the value of *α* that minimized the sum of squares. Extended Data Figure 1 illustrates this process given mean estimates of *Ae. aegypti* occurrence probabilities, *μ*(*T*), and *n*(*T*).

#### Attack rates under a statistical formulation

As an alternative to our model-based characterization of attack rates, we also considered a purely statistical approach that modeled probit-transformed seroprevalence observations as functions of averaged monthly temperatures, *Ae. aegypti* occurrence probabilities, and the economic index. We considered all combinations of linear, quadratic, and pairwise interaction terms of these variables, comparing them on the basis of Akaike Information Criterion using the lm and step functions in R^48^. Although additional functional forms would have been of interest, this suite of models was as complex as the limited set of 13 seroprevalence observations would support.

#### Quantifying uncertainty around attack rate projections

To quantify uncertainty associated with our projections, we generated 1,000 Monte Carlo samples from the uncertainty distributions of each model parameter. For *μ*(*T*) and *n*(*T*), we took random draws of their parameters consistent with published descriptions of uncertainty in the parameters of these functions from their original sources^17,18^. For *Ae. aegypti* occurrence probabilities, we drew randomly with replacement from 100 sample layers from the posterior distribution^7^. For the relationship involving the economic index and the *R*_0_ scaling factor *α*, we used best-fit SCAM models and *α* values corresponding to each set of random draws of the parameters of *μ*(*T*), *n*(*T*), and the *Ae. aegypti* layers. For each of the 1,000 Monte Carlo samples of the statistical model, we performed resampling with replacement among the 13 seroprevalence values, performed the same model fitting and model selection procedure described in the previous section, and took a multivariate normal random sample of the parameter values of the best-fit model based on the model’s best-fit parameters and variance-covariance matrix.

### Projecting attack rates and numbers of infections

To obtain estimates of numbers of infections in total and among childbearing women for the model-based and statistical approaches, we multiplied their respective attack rate projections applied to 5x5 km grids across Latin America and the Caribbean by human demographic layers for total population and births in 2015. For both the model-based and statistical approaches, we performed these calculations and summed at the country level once for each of the 1,000 Monte Carlo samples that we produced. High-resolution spatial projections of attack rates and numbers of infected childbearing women under the model-based approach are presented in Extended Data Figs. 2-10. Most projections based on the statistical approach resulted in attack rates of 100% in nearly all locations throughout Latin America and the Caribbean.

## ACKNOWLEDGEMENTS

We thank J Ashander, CM Barker, MA Johansson, RC Reiner, ST Stoddard, and members of the Perkins Lab for useful discussions. TAP, ASS, and AJT are supported by funding from NSF (DEB 1641130). TAP is supported by funding from NIH/NIAID (1P01AI098670-01A1) and the Bill & Melinda Gates Foundation (OPP1110495). AJT is supported by funding from NIH/NIAID (U19AI089674), the BMGF (OPP1106427, 1032350), NORAD, and a Wellcome Trust Sustaining Health Grant (106866/Z/15/Z). CWR is supported by funding through the University of Southampton’s Economic and Social Research Council’s Doctoral Training Centre. AJT and CWR acknowledge the support of the WorldPop (www.worldpop.org) and Flowminder Foundation (www.flowminder.org) teams in demographic dataset production, and TAP and ASS acknowledge support from the Notre Dame Center for Research Computing.

## EXTENDED DATA

**Extended Data Table E1.**
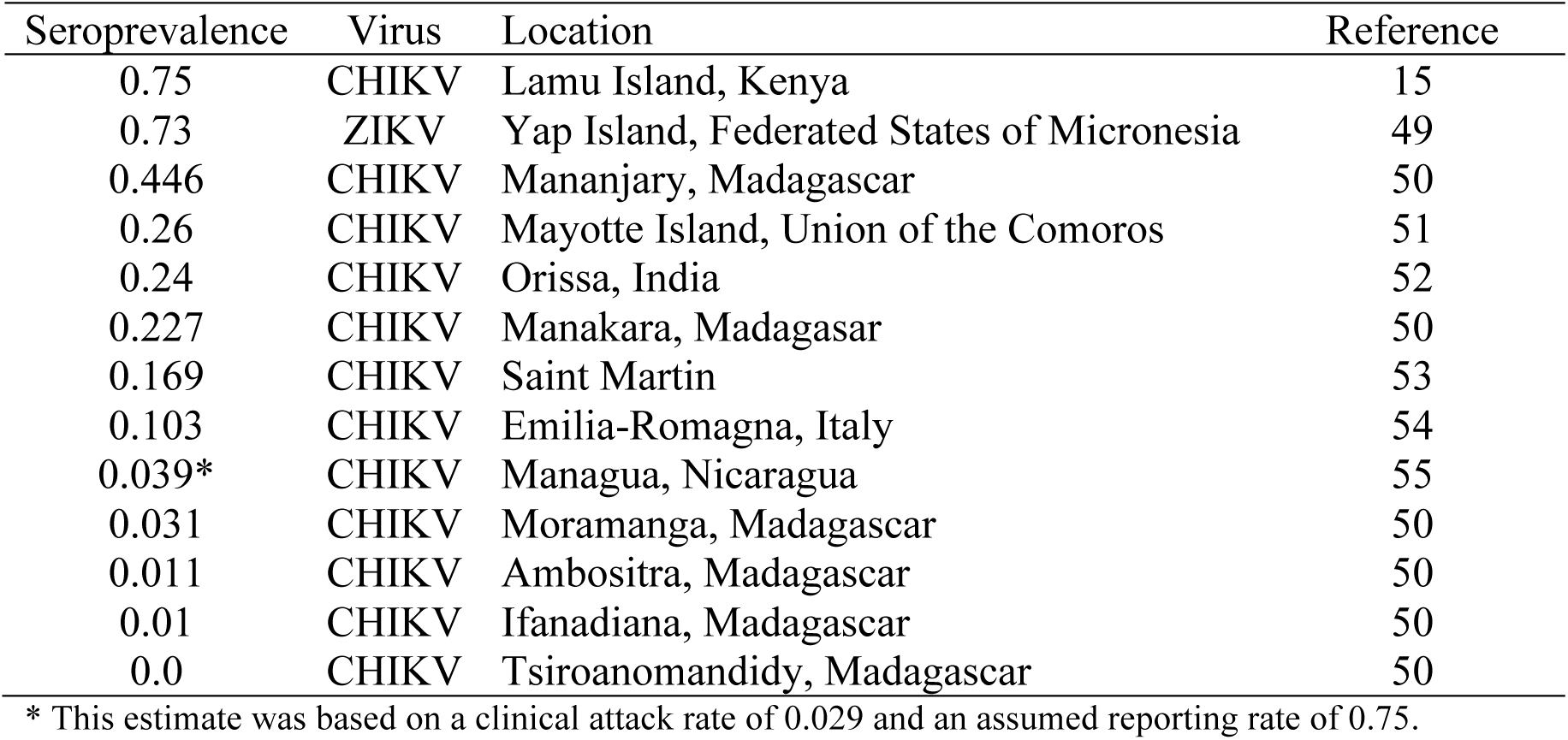
Seroprevalence estimates for Zika and chikungunya viruses in the context of recent oubtreaks in a previously susceptible population.

**Extended Data Figure E1.**
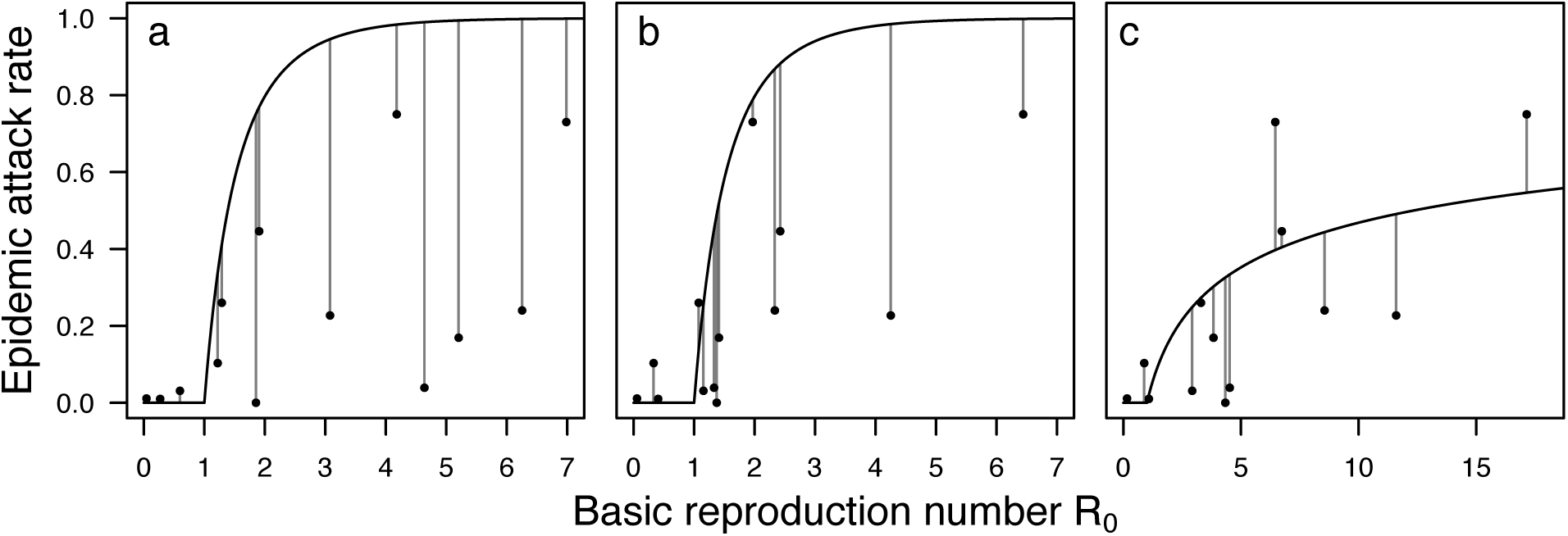
Relationships between the basic reproduction number *R*_0_ (x-axis) and epidemic attack rates (y-axis) under different model assumptions (a-c). In each panel, the curve shows this relationship under a given set of model assumptions and the points represent the 13 seroprevalence estimates (ED Table 1) mapped on to the *R*_0_-axis based on conditions at the sites where those data were collected. (a) *R*_0_ calculations according to eqn. 1 under the assumption that *m* = 1 and with parameter *α* = 1 determining the shape of the curve. (b) *R*_0_ calculations according to eqn. 1 under the assumption that *m* is determined by a SCAM model of the economic index and with parameter *α* = 1. (c) Same as b but with with parameter *α* = 0.13 fitted by least squares and a separate SCAM model fitted conditional on *α*

**Extended Data Figure E2.**
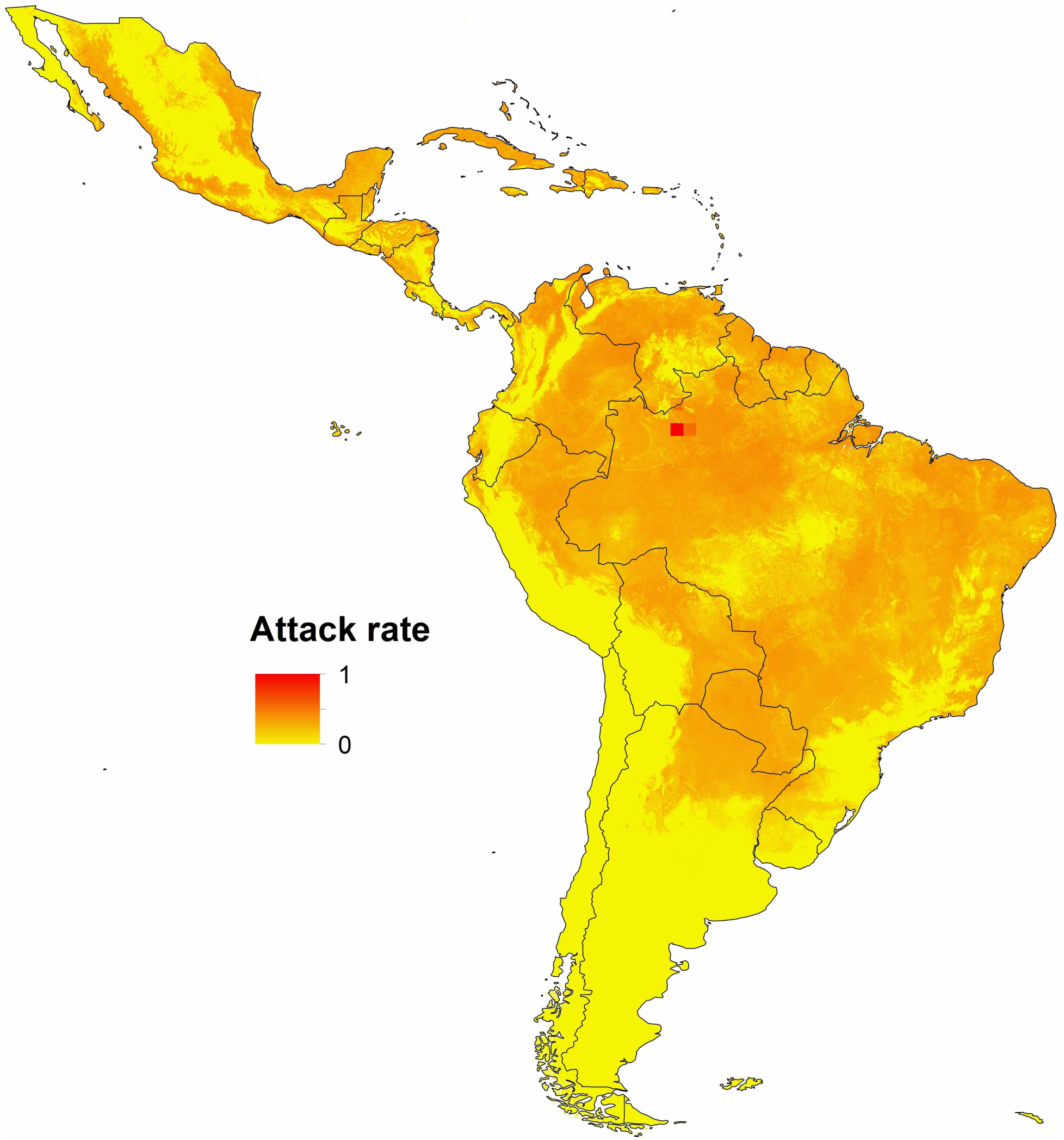
Gridded spatial projections of median epidemic attack rates at 5x5 km resolution in Latin America and the Caribbean. Each grid cell is shaded according to the median epidemic attack rate for that grid cell from the distribution of 1,000 Monte Carlo samples.

**Extended Data Figure E3.**
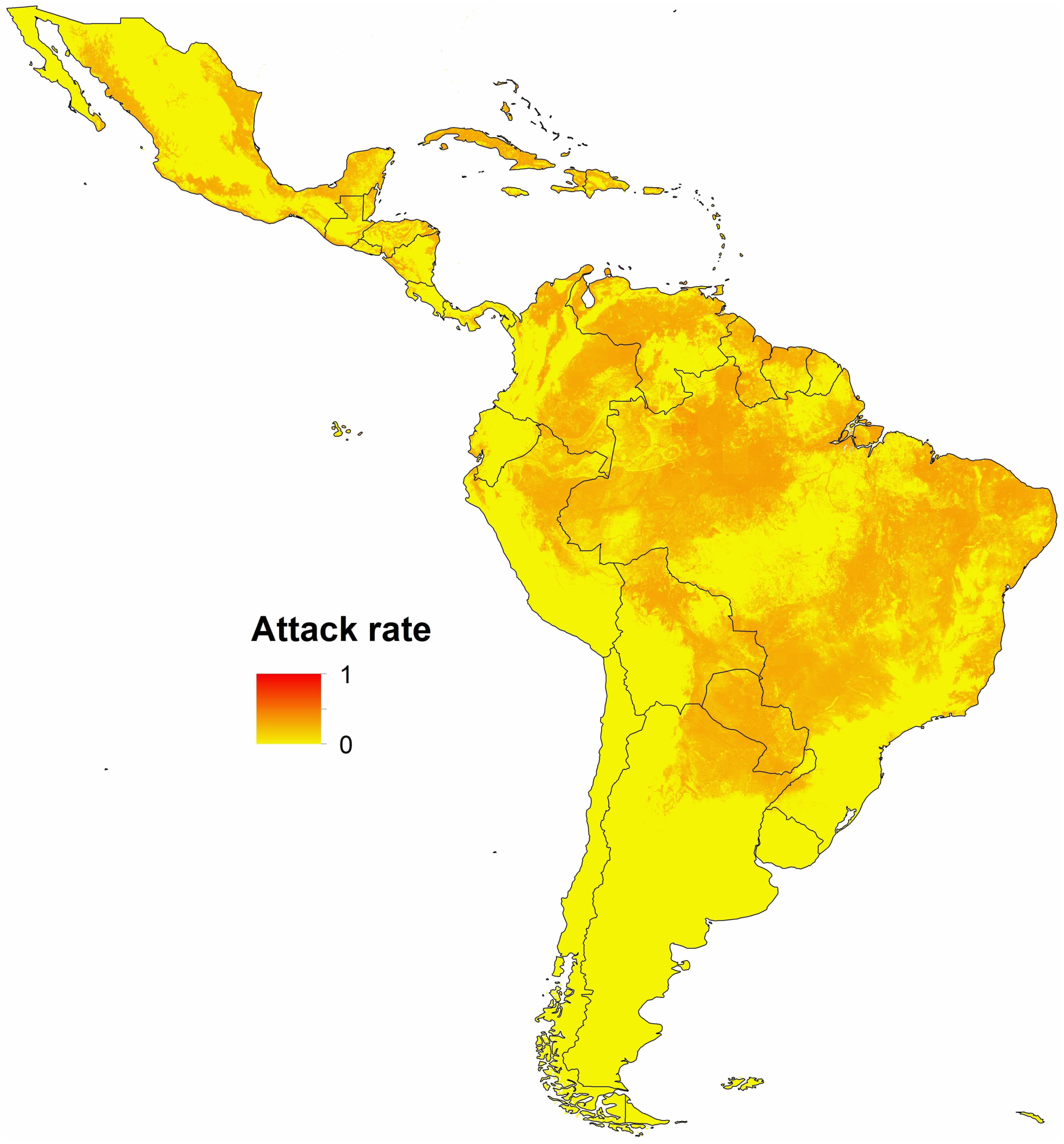
Gridded spatial projections of minimum epidemic attack rates at 5x5 km resolution in Latin America and the Caribbean. Each grid cell is shaded according to the minimum epidemic attack rate for that grid cell from the distribution of 1,000 Monte Carlo samples.

**Extended Data Figure E4.**
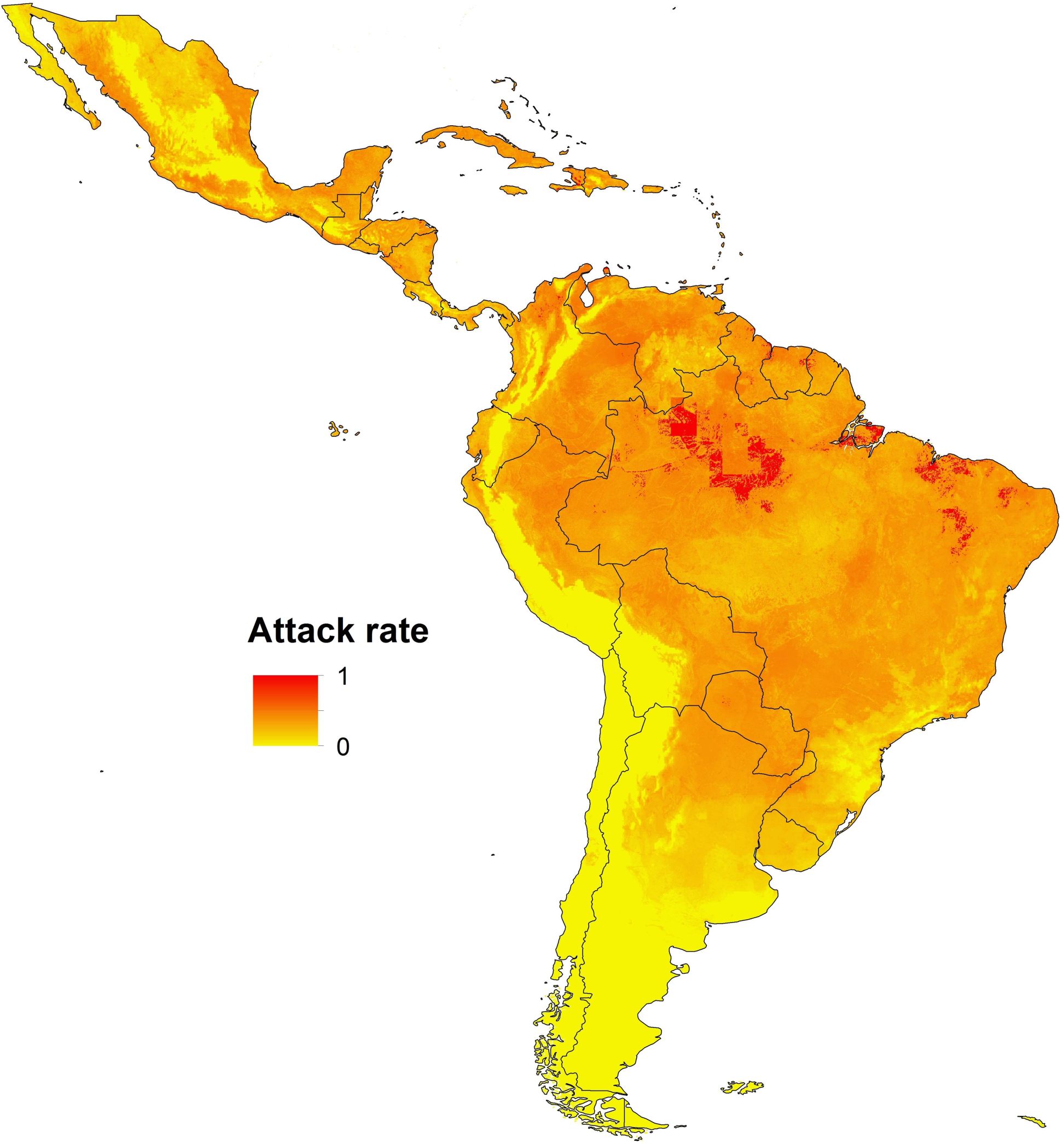
Gridded spatial projections of maximum epidemic attack rates at 5x5 km resolution in Latin America and the Caribbean. Each grid cell is shaded according to the maximum epidemic attack rate for that grid cell from the distribution of 1,000 Monte Carlo samples.

**Extended Data Figure E5.**
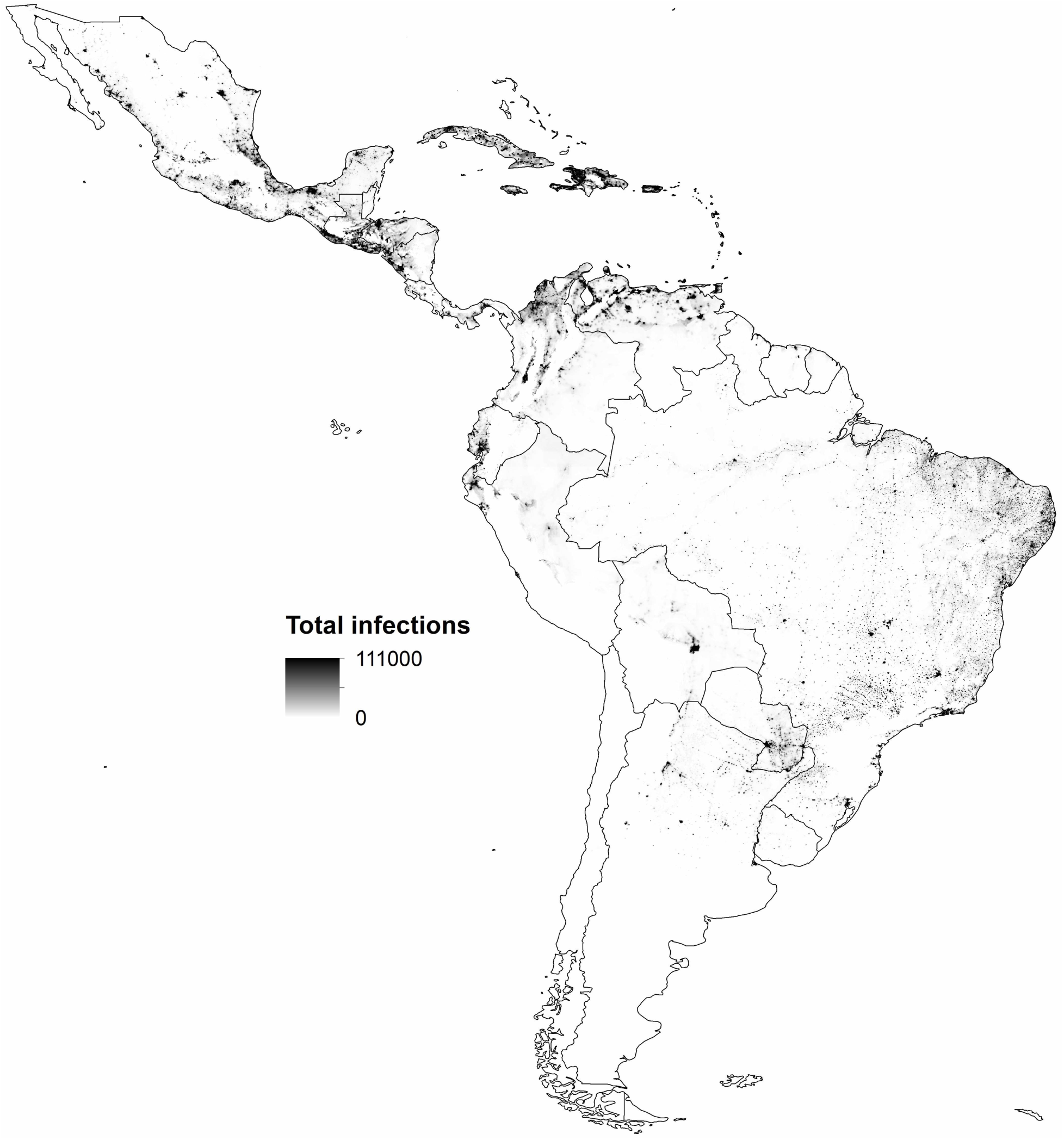
Gridded spatial projections of median infections among the total population at 5x5 km resolution in Latin America and the Caribbean. Each grid cell is shaded according to the median number of infections for that grid cell from the distribution of 1,000 Monte Carlo samples.

**Extended Data Figure E6.**
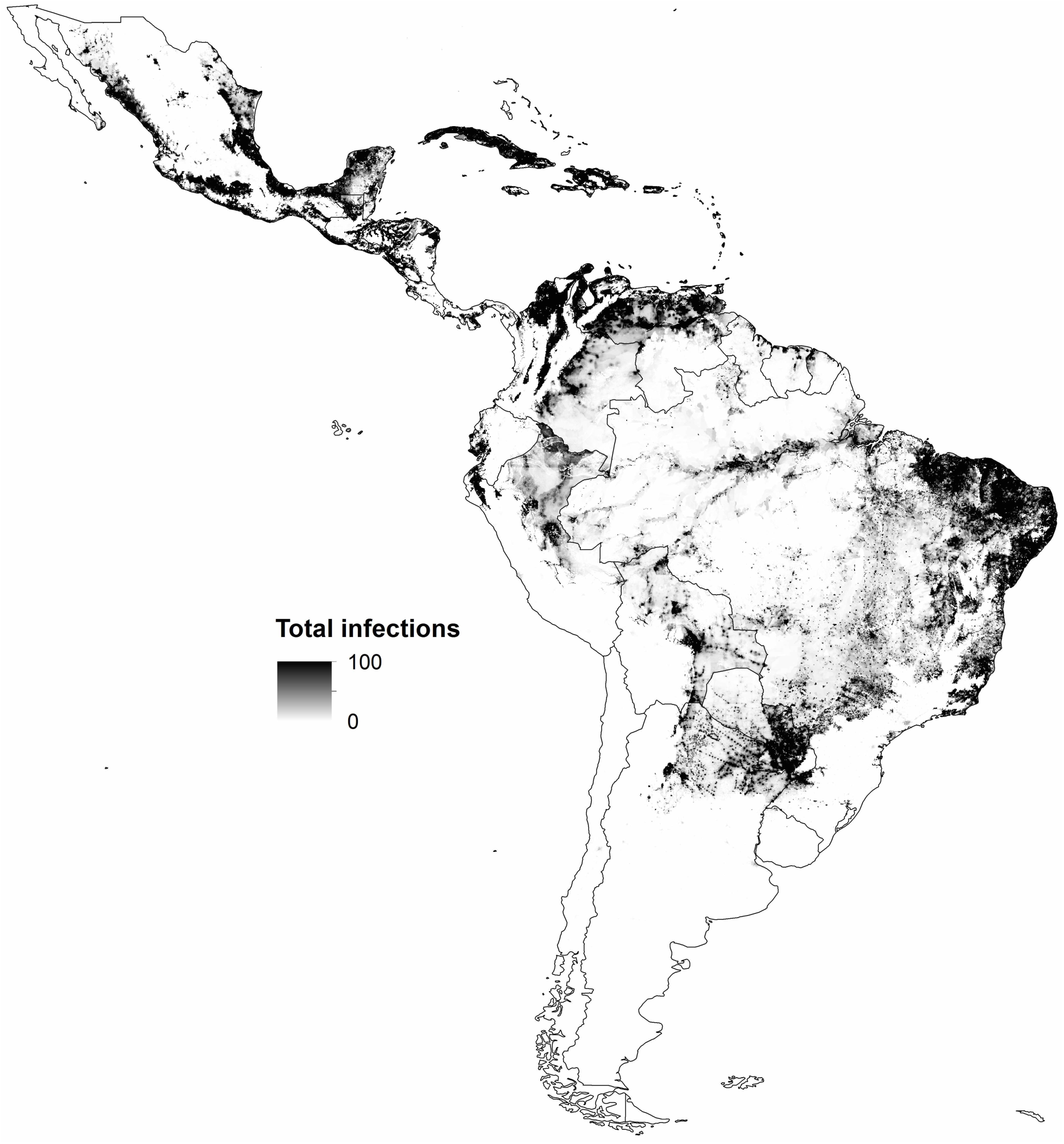
Gridded spatial projections of minimum infections among the total population at 5x5 km resolution in Latin America and the Caribbean. Each grid cell is shaded according to the minimum number of infections for that grid cell from the distribution of 1,000 Monte Carlo samples.

**Extended Data Figure E7.**
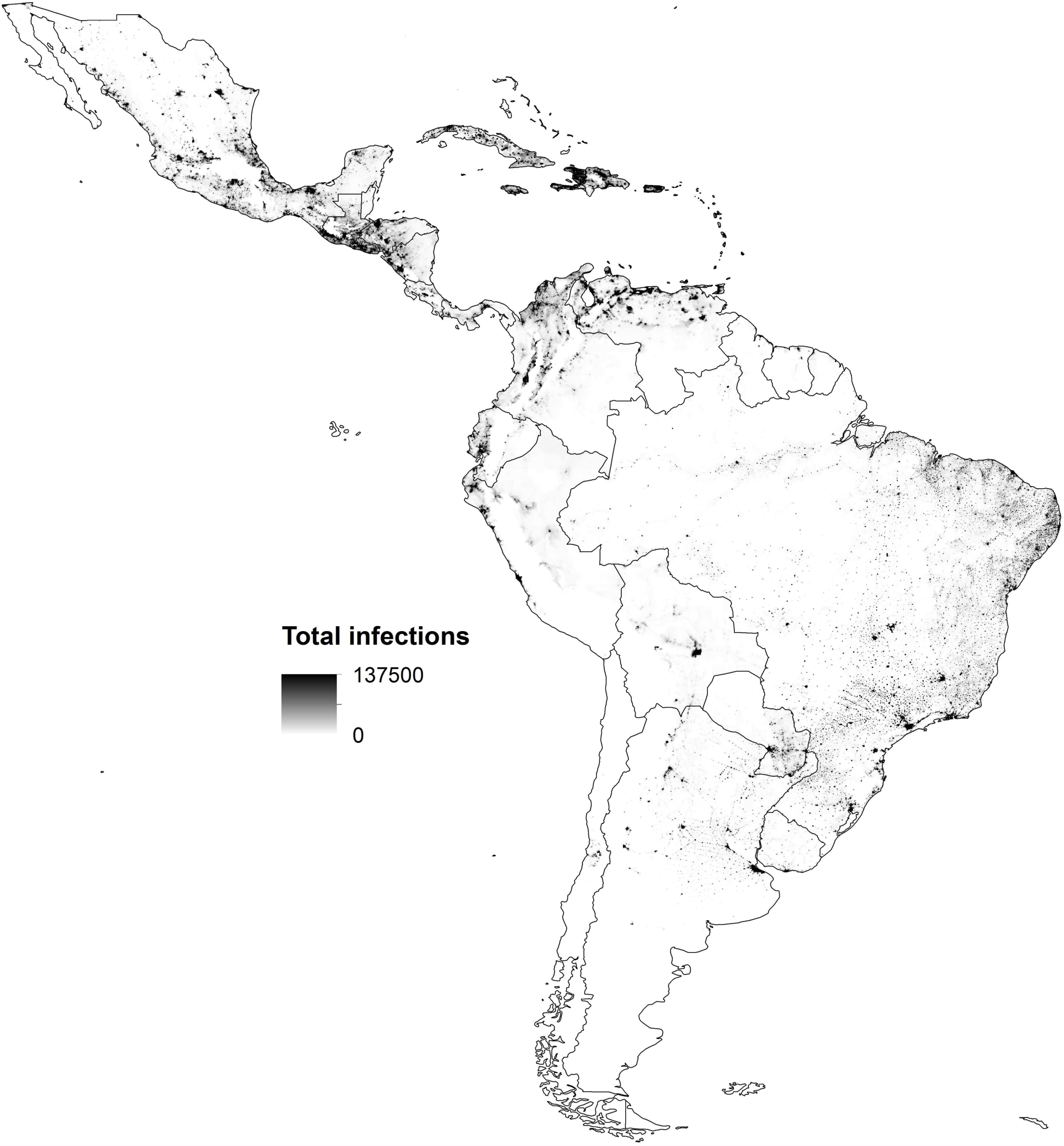
Gridded spatial projections of maximum infections among the total population at 5x5 km resolution in Latin America and the Caribbean. Each grid cell is shaded according to the maximum number of infections for that grid cell from the distribution of 1,000 Monte Carlo samples.

**Extended Data Figure E8.**
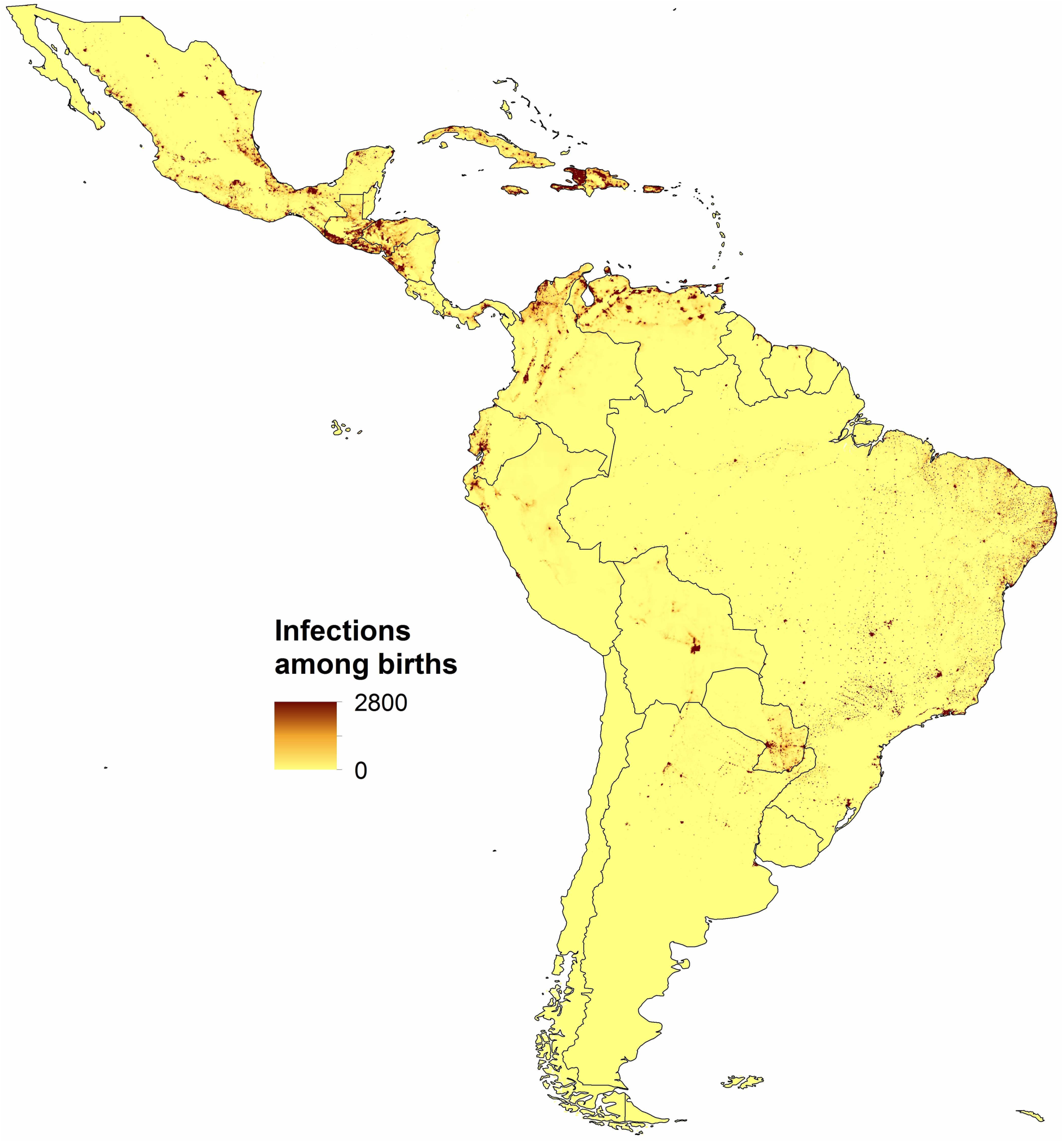
Gridded spatial projections of median infections among childbearing women at 5x5 km resolution in Latin America and the Caribbean. Each grid cell is shaded according to the median number of infections for that grid cell from the distribution of 1,000 Monte Carlo samples.

**Extended Data Figure E9.**
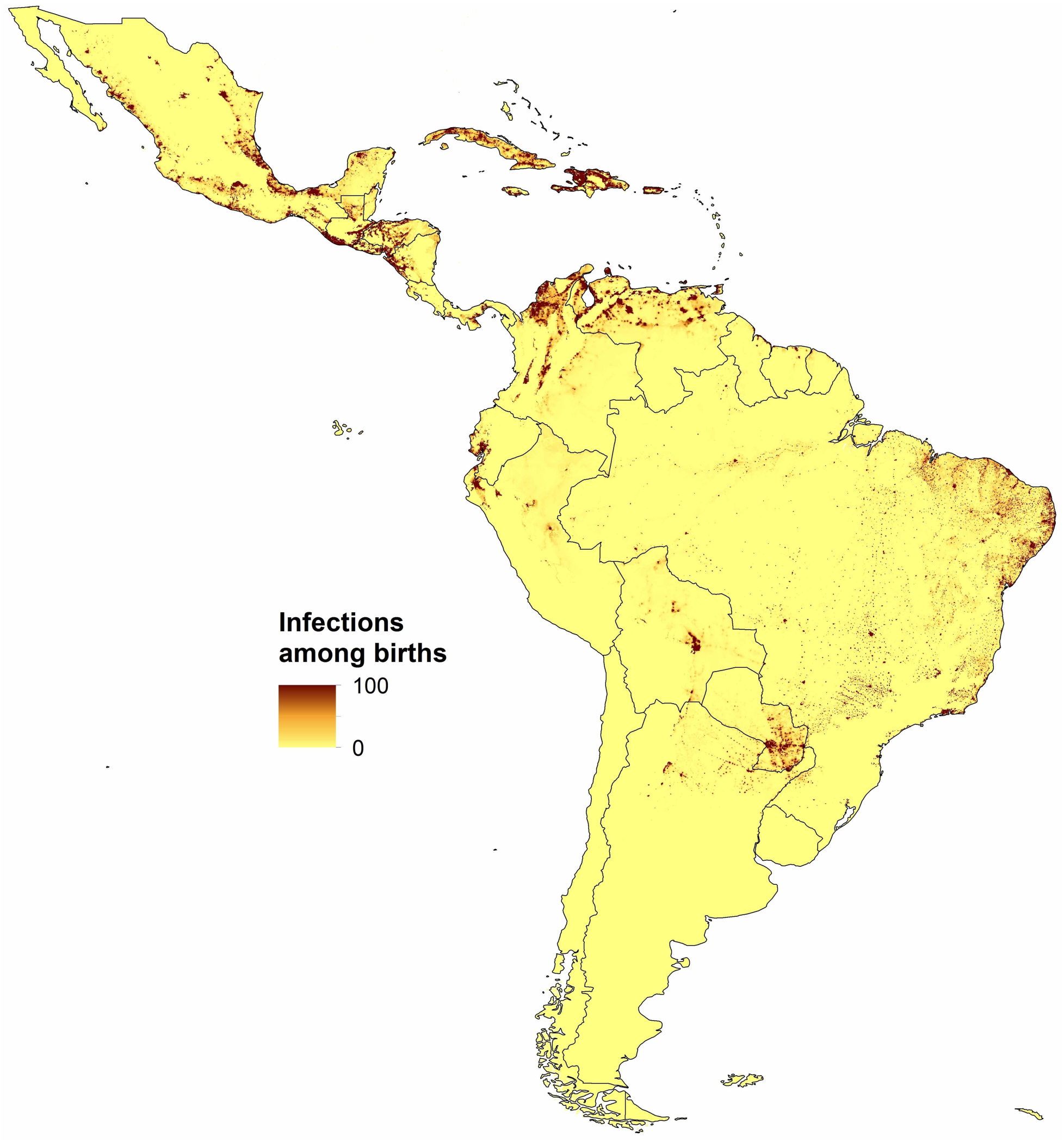
Gridded spatial projections of minimum infections among childbearing women at 5x5 km resolution in Latin America and the Caribbean. Each grid cell is shaded according to the minimum number of infections for that grid cell from the distribution of 1,0Monte Carlo samples.

**Extended Data Figure E10.**
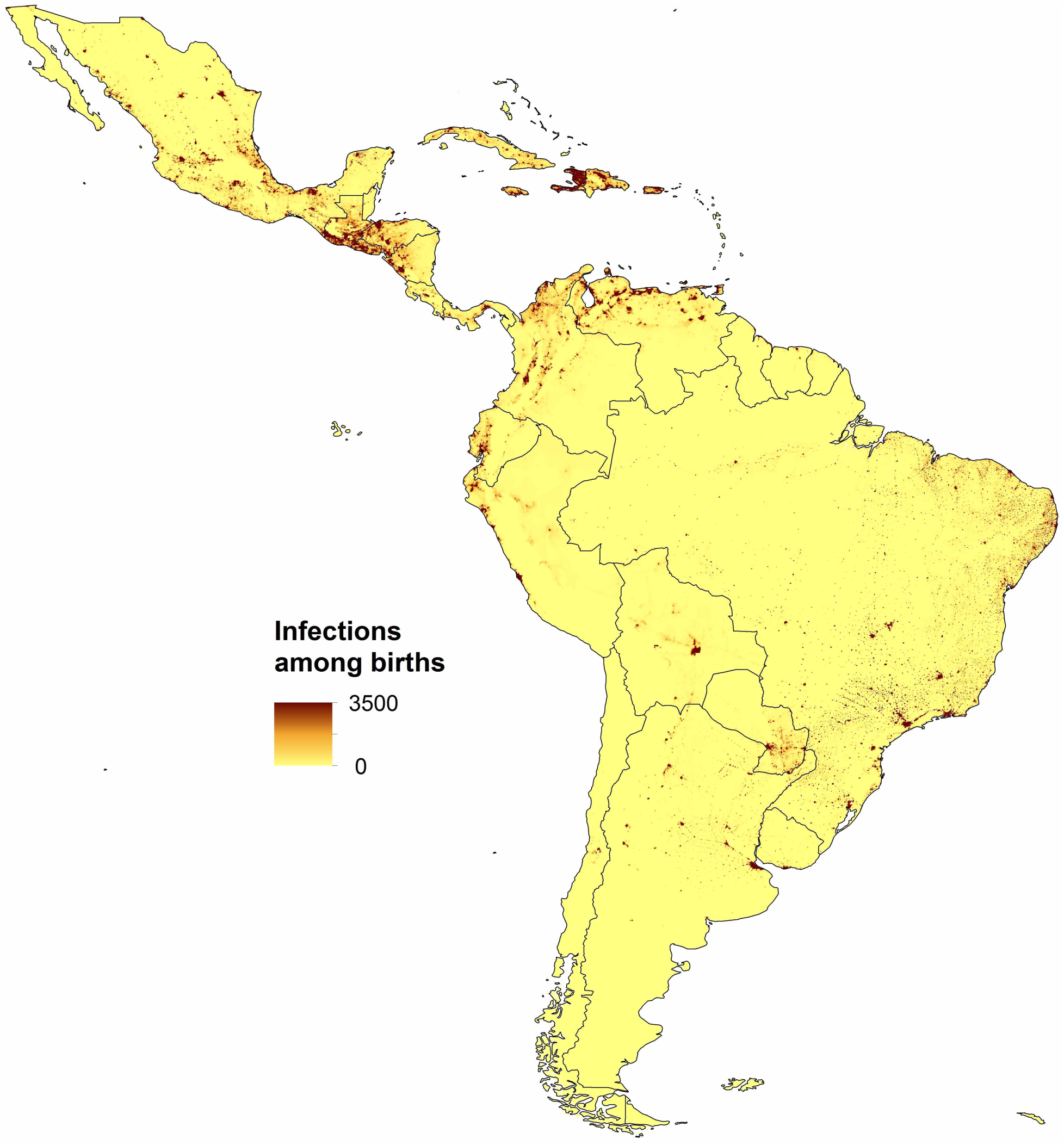
Gridded spatial projections of maximum infections among childbearing women at 5x5 km resolution in Latin America and the Caribbean. Each grid cell is shaded according to the maximum number of infections for that grid cell from the distribution of 1,0Monte Carlo samples.

